# Implications of (co)evolution of agriculture and resource foraging for the maintenance of species diversity and community structure

**DOI:** 10.1101/2021.03.02.433551

**Authors:** Aurore Picot, Thibaud Monnin, Nicolas Loeuille

## Abstract

Agriculture is found in numerous taxa such as humans, ants, beetles, fishes and even bacteria. This type of niche construction has evolved independently from hunting, though many species remain primarily predators. When a consumer has a positive effect on its resource, we can expect an allocative cost of agriculture, as the agricultural care diverts time and energy from other activities. Defending the resource against predators may divert time from its consumption (exploitation cost). The cost may also occur on the foraging of alternative resources, for instance if the consumer spends more time nearby the farmed resource and underexploiting resources elsewhere (opportunity cost). We here investigate transitions from predation to agriculture in a simple three-species model of a farmer that consumes two resources and has a positive effect on one. We study the conditions for the (co)evolution of the investment into agriculture and specialization on the two resources, and its consequences on the ecological dynamics of the community. Eco-evolutionary dynamics generate a feedback between the evolution of agriculture and specialization on the helped resource, that can lead to varying selected intensity of agriculture, from generalist strategies with no agriculture, to specialist farmers, with possible coexistence between these two extreme strategies.

## Introduction

Some of the species that display the greatest ecological success perform agriculture, that we here define as the cultivation of plants, algae, fungi and the herding of animal. Agriculture is a type of niche construction, an active modification of its environment by an organism, that potentially feed backs on the selective pressures acting upon it (Odling-Smee et al. 2013). A typical example is the impact of agriculture on the evolutionary history of humans and species affected by agriculture (Boivin et al. 2016). Humans have been causing both ecological and evolutionary change through their practice of agriculture, from direct effects such as artificial selection and domestication, to indirect effects such as climate change or introduction of invasive species, and causing the evolution of resistance to use of herbicides or pesticides (Thrall et al. 2011; Loeuille et al. 2013). The consequences of domestication, for instance, are important not only for the selected species (that can display drastic modifications of their phenotypic traits compared to the ancestral, undomesticated species (Thrall et al. 2011)) but also for other species within agricultural landscapes, as the domesticated species may become a dominant species exerting a strong selective force. Humans are not the only species to practice agriculture. Many ant species can use other insects, such as aphids, as cattle, or cultivate fungi (Mueller et al. 2005). Numerous taxa benefit from actively managing their resources: agriculture is also found in termites, beetles, fishes, nematodes and even microorganisms (Hata and Kato 2006; Boomsma 2011; Thutupalli et al. 2017; Brooker et al. 2020). The benefits associated with agriculture are easily understandable in changing, unpredictable and competitive environments: it potentially increases the resource availability, limits competition if the cattle or exploitation is privatized, and allows a greater predictability of resource abundance compared to foraging.

From a consumer-resource theory perspective, agriculture can be envisioned as a modification of a purely trophic interaction between a consumer and a resource. This interaction is then not only consisting of consumption or predation, but contains an additional positive effect of the consumer on the resource, which has been described in various contexts (Abrams 1992; Brown et al. 2004; Terry et al. 2017). The ecological and evolutionary consequences of including positive effects into trophic networks have recently received increasing attention (Fontaine et al. 2011; Kéfi et al. 2012; Mougi and Kondoh 2012). Accounting for such non trophic interactions alters the stability of networks and coexistence of species. Positive effects associated with agriculture can emerge from different consumer behaviours (protection against predators, helped reproduction or dispersal, for instance). It can impact the resource demography in various ways (increase of the carrying capacity, increase of the growth rate) that are expected to increase the resource profitability for the farmer species, compared to alternative foraged resources. The evolution of agriculture has been envisioned in the niche construction perspective, particularly in the human context (Rowley-Conwy and Layton 2011; Boivin et al. 2016; Zeder 2016) A classical example of the consequences of niche construction through agriculture is the evolution of lactose tolerance in humans (Laland and O’Brien 2011).

Jointly to the niche construction perspective, the ecological and evolutionary consequences of agriculture can be conceived in the light of foraging theories (optimal foraging, (Charnov 1976; Pyke et al. 1977), adaptive foraging, (Loeuille 2010)). This can help understand the question of the potential transition between foraging strategies such as hunting, gathering, to agricultural strategies. The evolution of agriculture can lead to full specialization and reciprocal dependency of the agricultural partners, as in fungus-growing ants (Mueller et al. 2005). To understand those two questions (transition to settled agriculture, extreme specialization), we can use the evolution of specialization framework (Egas et al. 2004; Abrams 2006). Optimal foraging theory predicts that the specialization of a consumer will evolve depending on the resources profitabilities. Agriculture practice can then modify the resource profitability through the modification of its abundance. Hence we expect that increasing agriculture selects for higher specialization on the cultivated resource that becomes more profitable, in turn selecting for more agriculture, leading to a positive feedback loop between agriculture and specialization on the cultivated resource. As a consequence, we expect reciprocal evolutionary consequences (or coevolution) between the evolution of specialization (*sensu* consumption, predation) on the resources, and the evolution of agriculture. From this perspective, we can understand the evolution of agriculture as the addition of another dimension to the classic evolution of specialization: an organism allocates energy or time between the consumption of different resources, and the cultivation of one or several resources. Two types of evolutionary outcomes are expected: either a high level of agriculture with a high specialization on the managed resource, or no agriculture and generalism or a specialist on another resource depending on the trade-off associated with the foraging activities.

This simple prediction can be modified by accounting for the cost of agriculture: the niche constructing phenotype can first be threatened by cheaters, if niche construction benefits are shared by all the population (potentially leading to a tragedy of commons, (Hardin 1968)). Farming can then be considered a public good (Thutupalli et al. 2017). Although privatization might overcome the threat of freerider invasion (through pleiotropy, (Chisholm et al. 2018), monopolization of the niche (Krakauer et al. 2009) or benefits going to the closest relatives (Scheiner et al. 2021)), trade-offs can still mitigate the evolution of agriculture by altering the profitabilities of the resources. If the cultivated resource is initially much less profitable than an alternative resource, we predict that agriculture may be counter-selected despite potential benefits. A cost to agricultural behaviour could emerge because of a high presence of competitors, predators or pathogens of the resource that needs to be actively protected (Hübner and Völkl 1996; Stadler and Dixon 2005; Adams et al. 2013; Fernández-Marín et al. 2013; Thutupalli et al. 2017). Here, we investigate the cost of agriculture in the foraging theory perspective: increasing the niche construction activity occurs at a cost on the consumption of resources, because of a limited energy or time budget available to the farmer species. Agriculture can, as a type of resource exploitation, decrease the consumption of an alternative resource (“opportunity cost”, described in Picot et al. (2019)). It can as well decrease the consumption of the managed resource, e.g. if defending it against predators or competitors implies moving away from the resource site, or allocating more time to defense rather than consumption (“exploitation cost” scenario, Picot et al. (2019)).

Because of the previously stated feedback loop, understanding the (co)evolution of agriculture and specialization requires to investigate its ecological consequences in terms of variations in resource abundances. Explicitly accounting for niche construction in models and theory leads to patterns that would not be predictable otherwise (Kylafis and Loreau 2011). In humans, agriculture is known to have drastic effects on the community and ecosystem level properties: for instance, by introducing invasive species that engage in apparent competition with native ones (David et al. 2017; Geslin et al. 2017), so that agricultural has pervasive consequences for the maintenance of species diversity and ecosystem functioning (Emmerson et al. 2016). The study of indirect effects may give insights on the community consequences of the evolution of agriculture. Indirect effects are effects of one species on another, transmitted by another species (Wootton 1994), here the consumer species. When a consumer consumes two resources, those resources engage in apparent competition (Holt 1977): an increase in density of one resource may decrease the other resource growth rate through their interaction with the consumer. Agricultural aspects modify this view. In an ecological analysis of a consumer-resource model with niche construction, Picot et al. (2019) show that cultivated and non-cultivated resources may then engage in various types of indirect interactions through the consumer. Without considering any cost, increasing niche construction has a positive effect on the managed resource density, which translates in a bottom-up positive effect on the consumer density, and a negative effect on the alternative density, because of apparent competition. Considering costs of niche construction mitigates this result. If the cost of niche construction is high enough in terms of resource consumption, an increase in niche construction of the cultivated resource may lead to counter-intuitive increase in the alternative resource density, because of a decrease in its consumption rate (“opportunity cost” scenario) and/or a decrease of the consumer density (“exploitation cost” scenario and “opportunity cost”).

In this study, we use adaptive dynamics and numerical simulations to study the evolution and coevolution of a consumer niche constructing trait and its specialization on two resources (Geritz et al. 1998). We address the following questions: (1) How does a fixed foraging strategy impact the selected investment into agriculture? We predict that a high specialization on the managed resource favours the evolution of agriculture, while a high specialization on the alternative resource prevents it. (2) How does the profitability of the different resources impact the coevolution of the niche construction trait and the specialization on the resources? We predict a positive correlation between the evolution of agriculture and specialization on the cultivated resource. (3) How do the evolution of niche construction and its coevolution with specialization impact the coexistence of the resources and the functioning of the system? We then predict that a high selected level of agriculture leads to the exclusion of the alternative resource because of increased apparent competition, while counter-selection of niche construction should lead to coexistence of the resources.

## Model presentation

### Ecological dynamics

Our model is based on ordinary differential equations describing the ecological dynamics of the consumer and two resource species. In this simple model (eq (1) and Fig 1 *A*), the consumer species *C* interacts with the two resources *R*_*1*_ and *R*_*2*_. The resource *R*_*1*_ is managed through agriculture: we assume it receives a positive effect that increases its growth rate, but it is also consumed. The alternative resource *R*_*2*_ is only consumed:

**Figure 1:**
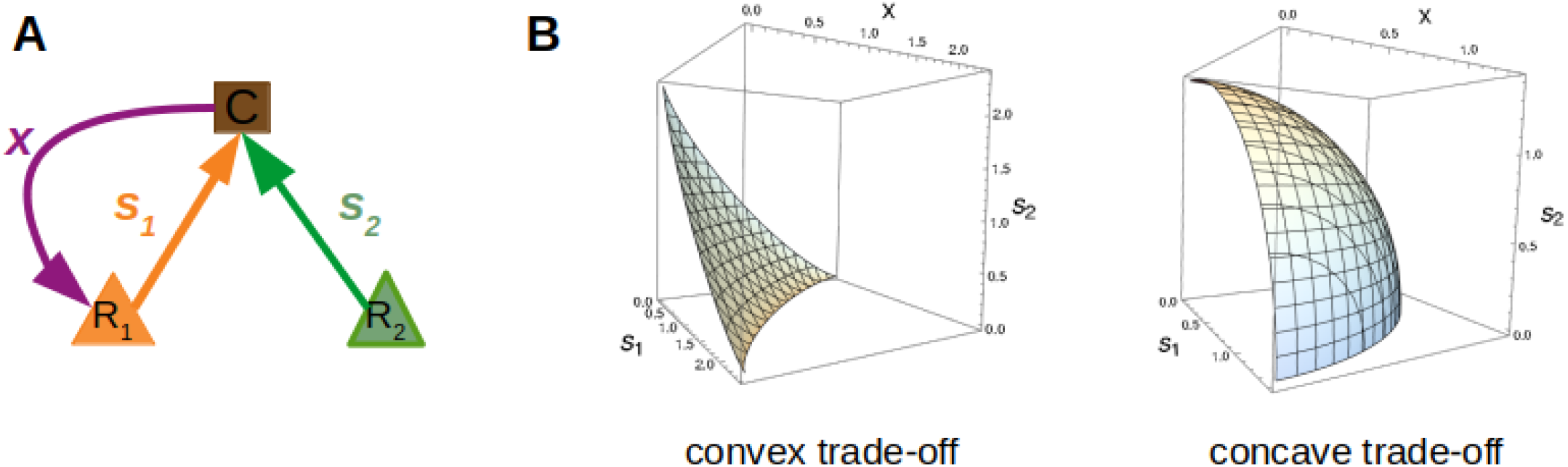
A) The module: the consumer *C* (in brown) consumes resources *R*_*1*_ (in orange) and *R*_*2*_ (in green) according to respective specialization rates *s*_*1*_ and *s*_*2*_. The resource *R*_*1*_ receives an additional positive effect through niche construction (agriculture) performed by the consumer. B) Different trade-off shapes determine the dependency between the three traits. Typically, concave trade-offs (here *k=2*) and convex trade-offs are considered (*k=0.8*).

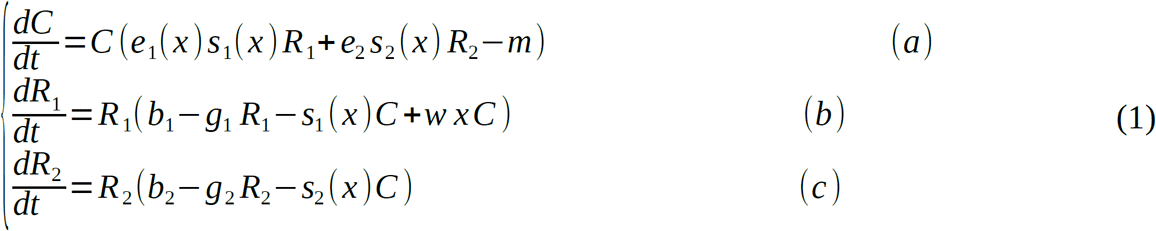

The consumer *per capita* growth rate depends on its consumption of resources (modelled by specialization on resource *R*_*i*_, *s*_*i*_, and consumption efficiency of resource *R*_*i*_, *e*_*i*_) and on its death rate *m*. In the absence of the consumer, the two resources grow logistically (allowing for their coexistence in such situations): *b*_*i*_ is the birth rate and *g*_*i*_ is the intraspecific competition rate for resource *R*_*i*_. Only apparent competition through the interactions with the consumer is considered here: we do not consider any direct competition between resources, which allows us to single out the impacts of the consumer. Finally, note that resource *R*_*1*_ receives a positive effect from agriculture, proportional to the investment into niche construction *x* and niche construction efficiency *w*. We assume that all parameters *e*_*i*_, *s*_*i*_, *b*_*i*_, *g*_*i*_, *x* are positive. Niche construction is then facultative for the maintenance of *R*_*1*_, as both resources have positive intrinsic growth rates.

### Trade-offs: costs and benefits of niche construction

As stated above, we assume that agriculture occurs at a cost for foraging on the resources. We consider two trade-off scenarios that rely on a time or energy constraint: increasing the agricultural intensity can decrease the consumption of the managed resource (“exploitation cost”, *s*_*1*_*’(x) < 0*) or the consumption of the alternative resource (“opportunity cost”, *s*_*2*_*’(x) < 0*). We consider that the three traits are linked through the following formula:

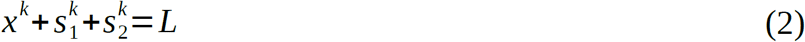

*L* represents the total amount of time or energy that the consumer allocates between niche construction and the consumption of the two resources. In the opportunity cost scenario, *s*_*1*_ is fixed while in the exploitation cost scenario, *s*_*2*_ is fixed. In the coevolution scenario, the three traits may evolve jointly along the trade-off surface (Fig 1 *B*). The exponent *k* shapes the trade-off (accelerating for a convex trade-off, *k* > 1 or saturating for a concave trade-off, *k* < 1).

We also assume that niche construction provides a direct benefit for the organisms performing it: for instance, agriculture can provide a more direct access to the managed resource and increase its consumption efficiency, because of proximity, or through another adaptation. We positively link the consumption efficiency to the agricultural trait through the following expression

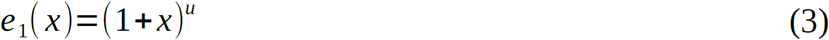

where *u<1* indicates saturating efficiency increase (diminishing returns, for instance if the efficiency consumption is constrained by other factors, such as physiological ones) while *u>1* indicates an accelerating response.

### Ecological dynamics

The ecological dynamics of a similar system has been studied in Picot et al. (2019), assuming a fixed level of niche construction, and differing only in the trade-off functions that are used. In this previous work, niche construction intensity linearly decreased with specialization rates *s*_*1*_ and *s*_*2*_, and no direct benefit of niche construction was assumed (this would mean *u=0* in our present model). However, some general results of the ecological dynamics apply in the present model: different types of ecological equilibria are obtained: either the coexistence of the two resources, with or without the consumer, or the maintenance of one resource (still with or without the consumer) or the extinction of all species. Note that this previous work only tackled ecological dynamics, while we here focus on the evolution of the different traits.

### Evolutionary dynamics

We first study the evolution of the phenotypic trait *x* using the adaptive dynamics framework (Dieckmann and Law 1996; Geritz et al. 1998), assuming that the consumer diet (*s*_*1*_,*s*_*2*_) is fixed. Adaptive dynamics allows to investigate evolutionary dynamics of phenotypic traits while defining the fitness of a given phenotype based on its ecological dynamics. The analytical framework relies on the separation of ecological and evolutionary timescales, while numerical simulations allows us to more freely investigate the limits of this hypothesis. The evolution of a trait is studied through several steps, assuming clonal reproduction, and small and rare mutations:

- the ecological equilibrium is determined, for a monomorphic population of resident trait *x*_*res*_ (by nullifying equations 1(a) to 1(c))
- a rare mutant with trait *x*_*mut*_ is introduced and replaces the resident trait if its invasion fitness (i.e., its *per capita* growth rate when rare, based on eq1(a)) is positive. Invasion fitness is given by:

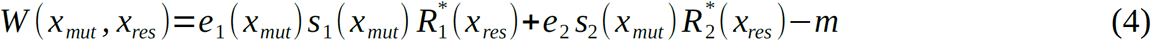

The new ecological equilibrium is established and the process is iterated.

A deterministic approximation of the evolution of the trait is given by the canonical equation of adaptive dynamics (Dieckmann and Law 1996) :

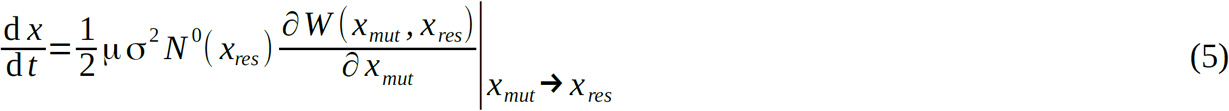

μσ^2^ *N*^0^(*x*_*res*_) is the evolutionary potential, that is total phenotypic variability brought by mutations with *μ* the *per capita* mutation rate, *σ*^*2*^ the phenotypic variance associated to these mutations and *N*^*0*^*(x*_*res*_) the resident population equilibrium density. The selection gradient 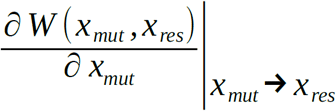 is derived from the invasion fitness (eq 4) and corresponds to the slope of the fitness landscape around the resident population, therefore representing the selective pressures acting on the phenotypic variability brought by mutations. The traits that nullify this gradient are called singular strategies (Dieckmann and Law 1996). Dynamics around these singular strategies are characterized by the second partial derivatives of fitness (Dieckmann and Law 1996; Geritz et al. 1998). This allows to distinguish two stability-related properties: invasibility and convergence. A singular strategy *x\** is non-invasible (Maynard Smith 1982), or evolutionary stable if:

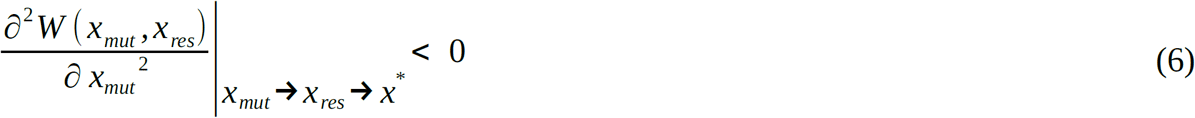

*i.e*. when the singular strategy is the resident population, no nearby mutant can invade.

A singular strategy is convergent or continuously stable (Eshel 1983) if:

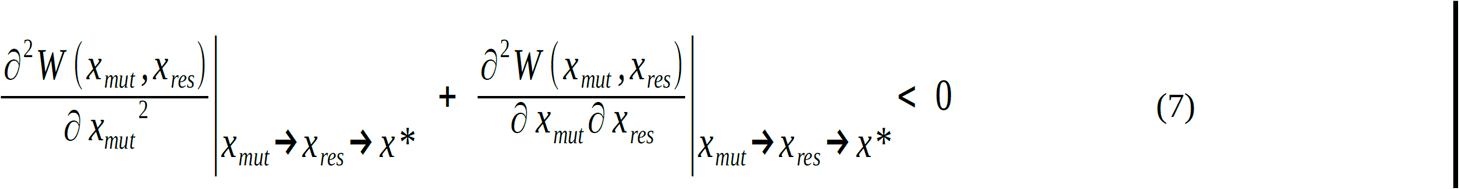

*i.e*., considering a resident population close to the singular strategy, mutants that are even closer to it are selected. The two trade-off scenarios lead to different fitness expressions for each scenario, since in the “exploitation cost” scenario, *s*_*2*_ is fixed and *s*_1_ 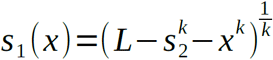 while in the opportunity cost scenario *s*_*1*_ is fixed leading to 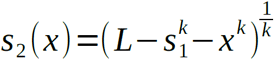.

We illustrate the evolution of *x* in the “exploitation cost” and “opportunity cost” scenarios using numerical simulations. The simulation algorithm was built in C++.

### Coevolutionary dynamics

In the coevolution scenario, we let the three traits *s*_*1*_, *s*_*2*_, and *x* vary while accounting for trade-off constraints (equation (2)). This means we allow the two traits *s*_*1*_ and *x* to evolve independently and derive the trait *s*_*2*_ from the trade-off. We approach this question both with semi-analytical calculations, and using our evolutionary algorithm, by introducing mutations of the three traits.

When both *x* and *s*_*1*_ jointly evolve, we derive two expressions of the fitness. The fitness *W*_*x*_*(x*_*mut*,_ *x*_*res*_, *s*_*1res*_) of a mutant of trait *x*_*mut*_ and the fitness *W*_*s1*_*(s*_*1mut*_, *s*_*1res*_, *x*_*res*_) of a mutant of trait *s*_*1mut*_ appearing in a resident population of traits *x*_*res*_ and *s*_*1res*_ are:

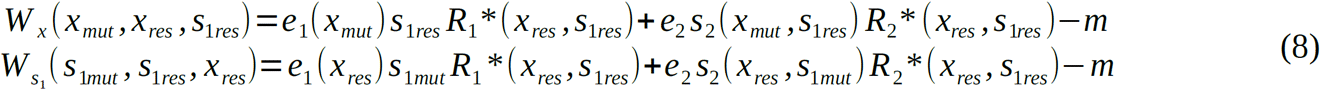

We can then compute the two coupled fitness gradients:

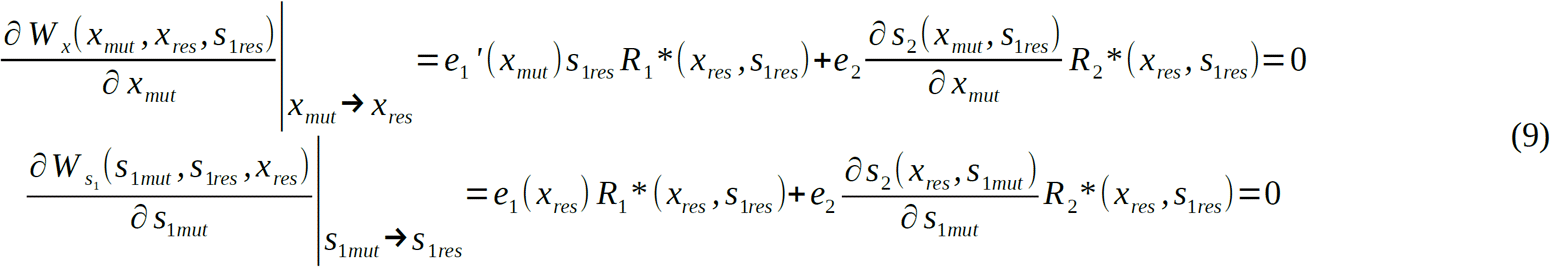

The coevolutionary dynamics of the traits are then described by a system of coupled canonical equations (Dieckmann and Law 1996):

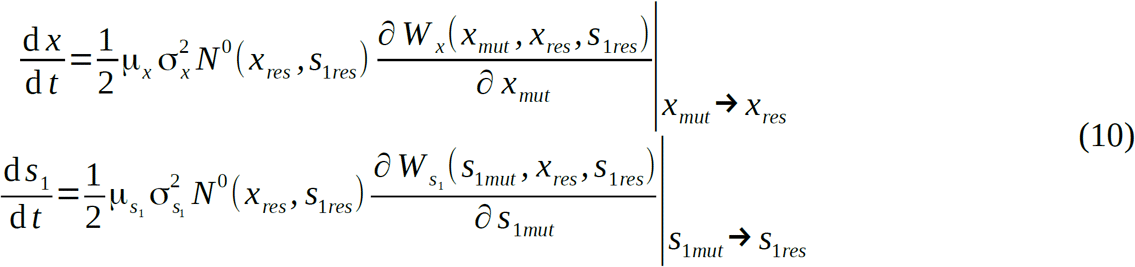

The singularities are obtained by solving the system (10), which means finding *x* and *s*_*1*_ values that simultaneously nullify the two fitness gradients, *(x*,s*_*1*_*\*)*. This is done by numerically solving the equations in Mathematica with fixed parameter values. From the equations of the coevolutionary dynamics, we can derive the Jacobian matrix and evaluate the conditions to obtain a stable coalition (evolutionary attractor) for the singularity *(x*,s*_*1*_*\*)* (Marrow et al. 1996).

The Jacobian matrix *J* of the system (10) at the equilibrium *(x*,s*_*1*_*\*)* is:

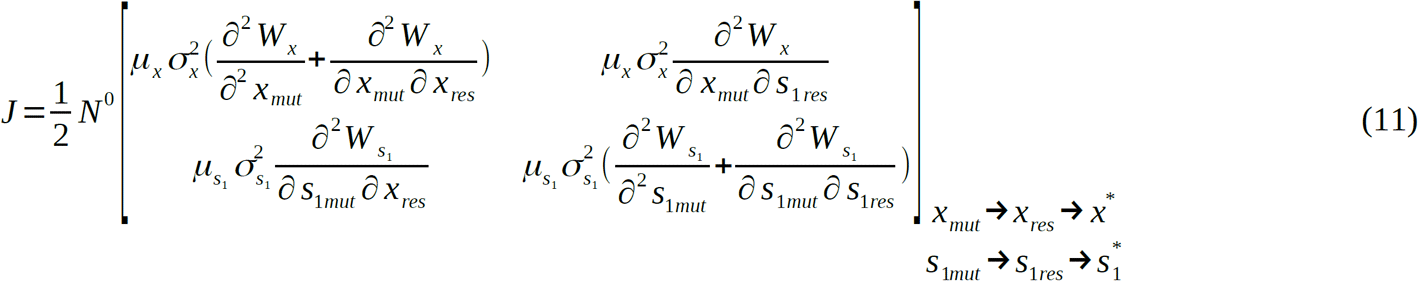

The condition to have a stable equilibrium *(x*,s*_*1*_*\*)* is that *Tr(J)* < 0 and *Det(J)* >0. We note that the diagonal terms correspond to the criteria of convergence in the monomorphic evolution, so if these terms are negative, then *Tr(J)*<0. The second condition is that *Det(J)*>0 (which is the condition for absolute convergence in Kisdi (2006). This condition can be expressed as:

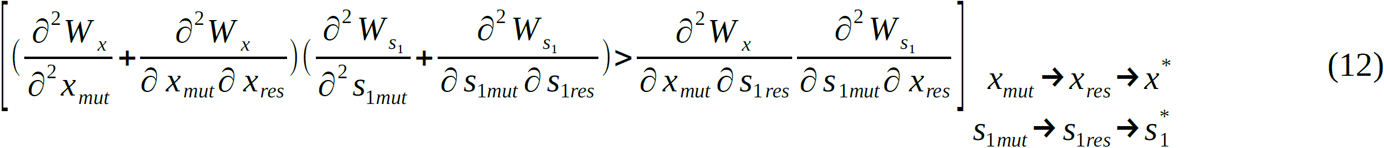

We then numerically check that the two conditions are met at the singularities. We also adapt our evolutionary algorithm in C++ to take account for the coevolution of the three traits and check that the algorithm converges to the analytically obtained singularity values.

## Results

### 1) Evolutionary dynamics of agriculture in the two trade-off scenarios

#### “Exploitation cost” scenario: managing the resource or consuming it

In this scenario, increasing the intensity of niche construction negatively impacts the consumption of the helped resource. We first study the general possible evolutionary dynamics depending on the trade-off shape, then focus on the linear trade-off scenario to get a more thorough mathematical investigation. As a first step, we do not specify the various functions implied in the biological trade-off, in order to have a general analysis of possible evolutionary dynamics: we ignore equations 2 and 3 and simply assume that *s*_*1*_ decreases with *x* while *e*_*1*_ increases with *x*.

Considering these assumptions in the fitness function definition (eq (4)) allows us to compute the fitness gradient (eq (13)):

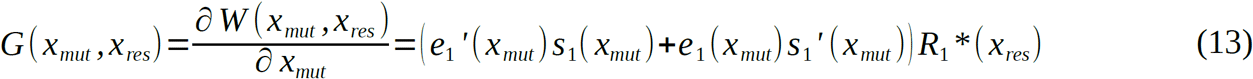

From this equation, we immediately note that if we do not consider a direct benefit to niche construction (*e*_*1*_*’(x*_*mut*_)=0), the fitness gradient becomes negative (as *s*_*1*_*’(x*_*mut*_)<0). This leads to gradual decrease of the trait and counter-selection of niche construction. This is consistent with the result of Chisholm et al. (2018): considering a direct benefit (from pleiotropy or pseudo-spatialization) is necessary to avoid a tragedy of the commons. That is inevitable given that our completely mixed model implicitly assumes that all consumers have access to the farmed resource (i.e., interaction is modeled by a mass action function). Variation in spatial access is a well-known way to avoid this situation (Lion et al. 2011). To keep the model simple and tractable, we thus consider an easier access to the constructed ressource, i.e.that *e*_*1*_*’(x*_*mut*_*)>0*.

Assuming that the helped resource is not extinct (*R*_*1*_ ^*\**^*(x*)* > 0), the position of singular strategies *x\** can be obtained from eq (14).

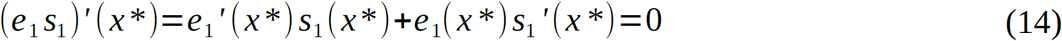

This mathematical condition means that the singular strategies correspond to an optimum of the realized consumption of the managed resource: if this strategy is an evolutionary endpoint, evolution optimize total effective consumption of the managed resource, since the consumption efficiency and specialization rate vary oppositely with agriculture trait *x*. We then expect intermediate levels of agriculture to be selected for. In the scenario, we also note that the value of the singular strategy only depends on this consumption-efficiency optimization: the densities of the resources do not matter.

Replacing the fitness gradient with the chosen functions of equations 2 and 3 (in equation (14)) leads to:

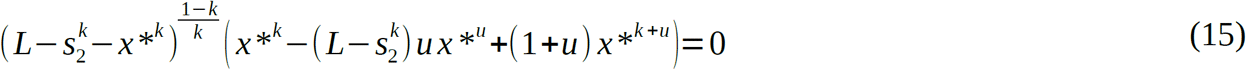

The condition for non-invasibility and convergence (that are equivalent here, see Supplementary Information for details) is:

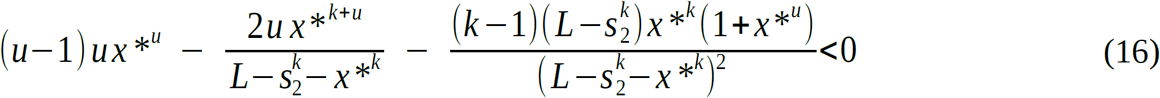

In the linear case (*u=k=1*), the equation (15) becomes (*x* *−(*L*−*s*_2_) *x* *+2 *x* *^2^)=0. We obtain two possible singularities: *x\** = 0 (which is a neutral case in the sense that the second derivative is null) or 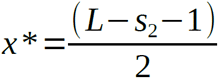 which is always convergent and non-invasible (CSS) (the second derivative expressed in (16) is 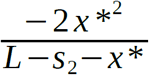 and is negative), it is an evolutionary endpoint.

While a completely general study using non-linear trade-offs is not possible, note that a sufficient condition to obtain a CSS is that *u* < 1 and *k* > 1 (see equation (16)), in other terms having diminishing returns of niche construction on resource consumption efficiency and a concave trade-off between specialization and niche construction. If *u* > 1 and *k* < 1 (accelerating efficiency and convex trade-off) the strategy may be characterized as a repellor (an unstable evolutionary point) depending of the sign of our expression. In this case, directional selection brings the trait to either zero niche construction (and total consumption), or total niche construction (and zero consumption), the latter case being less likely biologically. We show the possible evolutionary dynamics for different trade-off shapes in Supplementary Figure S1.

#### “Opportunity cost” scenario: managing one resource or foraging on another resource

When increasing agriculture occurs at a cost of the consumption of the alternative resource, the invasion fitness of a mutant of trait *x*_*mut*_ appearing in a resident population of consumer with trait *x*_*res*_ is derived:

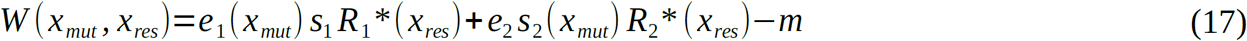

To obtain the fitness gradient *G(x*_*mut*_,*x*_*res*_) we consider the derivation of the fitness with respect to the mutant trait *x*_*mut*_:

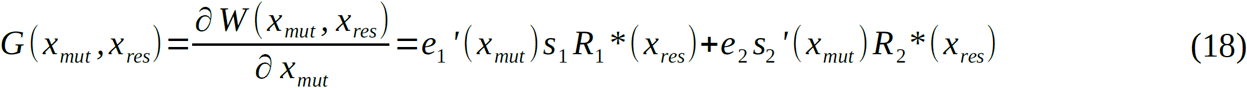

Singularities *x\** are obtained through nulling *G(x*,x*)*. The position of the singularity depends on optimizing both the consumption efficiency and the cost on the alternative resource consumption. Contrary to the previous scenario, the resident agricultural trait *x*_*res*_ matters for the position of the singularity. The non-invasibility condition is given for a general trade-off and for our chosen trade-off functions in equation (18):

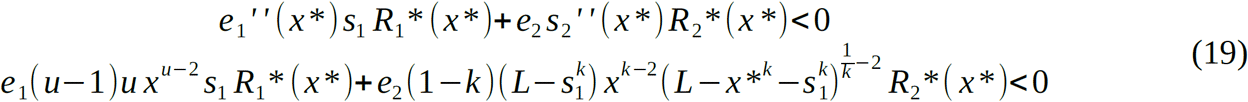

In the linear case (*u* = *k* = 1), the invasibility criteria is null, but not necessarily the convergence criteria. This can lead us to identify parameters set for which the strategy is convergent, and invasible.

As in the previous scenario, we can get partial information in the non-linear cases. We obtain a sufficient condition for non-invasibility of the singular strategy (equation (19)) (i.e. if this level of agriculture is reached, it is an ESS): *u < 1* and *k > 1*. A sufficient condition for the invasibility of the point is that if *u > 1* and *k < 1*. Those invasibility conditions are the same as for the “exploitation cost” scenario invasibility condition. Conditions for evolutionary convergence are however not tractable in the general case, and we only study numerically (see Supplementary Information for the convergence condition).

### 2) Effect of specialization patterns on the evolutionary dynamics of agriculture and feedbacks on the ecological dynamics

In the “exploitation cost” scenario, when we consider linear functions, we obtain a continuously stable strategy (CSS) that is both convergent and non-invasible. The value of this selected level of agriculture is expressed in eq 20:

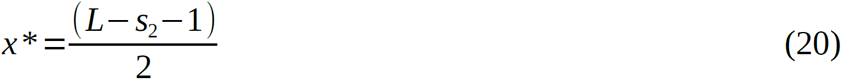

This singular strategy is positive if *s*_*2*_ *< L– 1*. If *s*_*2*_ *> L–1*, the value of the singularity becomes negative and the niche constructing trait evolves to 0. From the expression of the singularity (eq 20) we show that the selected level of niche construction is is negatively correlated to specialization on resource *R*_*2*_, consistent with our predictions.

The consumption of the managed resource at the evolutionary equilibrium (*s*_*1*_*(x*)*) is

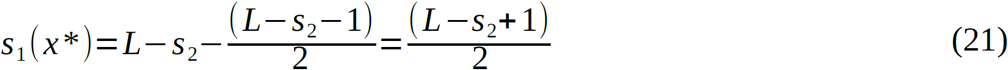

We plot the value of the selected agriculture level and consequent consumption of the managed resource, as a function of specialization on resource 2 in Figure 2 *A*. We represent the area of stable coexistence of the species when resource 2 specialization *s*_*2*_ and *x* vary (ecologically). We show that the selected value of agriculture, *x\** (represented as a black line) falls in the resource coexistence area (the black arrows indicated the direction of evolution, meaning that starting from a high value of agriculture, evolution brings the trait back into a coexistence area. The corresponding equilibrium densities are shown on Figure 2 *B*. Increasing the consumption of the alternative resource leads to a decrease in its evolutionary equilibrium density. The highest consumer equilibrium density is obtained when the specialization on the alternative resource is intermediate (and when the cultivated resource density is the lowest).

**Figure 2:**
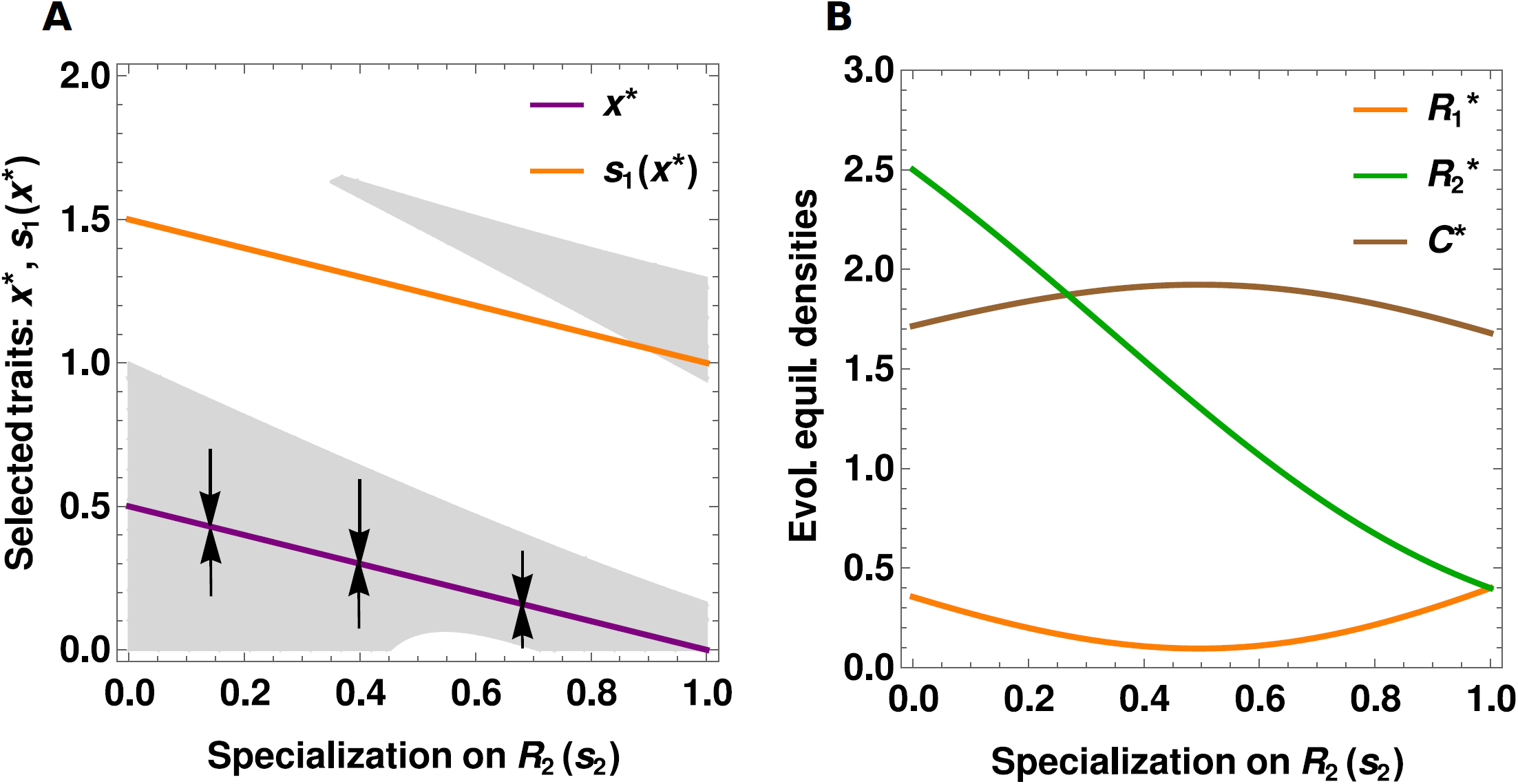
Effect of specialization on resource 2 on the evolutionary and ecological dynamics in the “exploitation cost” scenario. A) effect on the selected level of niche construction *x\** (in purple, the arrows indicate the direction of evolution) and subsequent consumption of resource 1, *s*_*1*_*(x*)* (in orange). The area of stable ecological coexistence of the three species when varying agriculture and the specialization on *R*_*2*_ (i.e., in a *(s*_*2*_,*x)* plane) is represented in grey. B) effect on the species densities at the ecological evolutionary equilibrium (that is at the selected niche construction): the consumer is in brown, *R*_*1*_ is in orange, *R*_*2*_ is in green. *b*_*1*_ *= b*_*2*_ *= 2, g*_***1***_ *= g*_*2*_ *= 0.8, e*_*1*_ *= e*_*2*_ *= 1, L =2, w = 1, m = 0.8, k=1, u=1*

The slopes of the selected *x\** and *s*_*1*_*(x*)* are the same (the lines are parallel), meaning that a higher specialization on resource 2 equally impact niche constructing activities and exploitation of the managed resource. This comes directly from our trade-off expression: mathematically, we assume that the total amount of energy or time *L* is a linear combination of all the traits and here they all have the same coefficient (i.e., 1). Were we to weight the different traits through different coefficients, so that the variation of each trait does not have the same impact on the energy allocation, then at the evolutionary equilibrium, modifying the specialization on resource 2 would lead to weighted cost on agriculture investment and the resource 1 consumption (see Supplementary Figure S2). In the non-linear CSS cases as well, the intensity of alternative resource consumption is negatively correlated with the evolved level of agriculture (figure not shown). This result is qualitatively the same when the strategy is a repellor. The singular strategy value is positively related to the specialization on *R*_*2*_, so that a higher specialization *s*_*2*_ on the alternative resource creates a larger basin of attraction for the zero niche construction state, i.e., niche construction is more often counter-selected.(see Supplementary Figure S3).

In the “opportunity cost” scenario, we do not obtain an analytical value of the expected selected level of agriculture: it is obtained by numerically nullifying the fitness gradient (equ. (18)), and then computing convergence and invasibility criteria. We do this for a range of *s*_*1*_ values in order to investigate our prediction that higher specialization on the managed resource should select for higher investment in agriculture. On Figure 3, we show the selected investment into agriculture in CSS cases (evolutionary endpoint), as well as the resulting specialization on the alternative resource, and the densities of the species at the evolutionary equilibrium.

**Figure 3:**
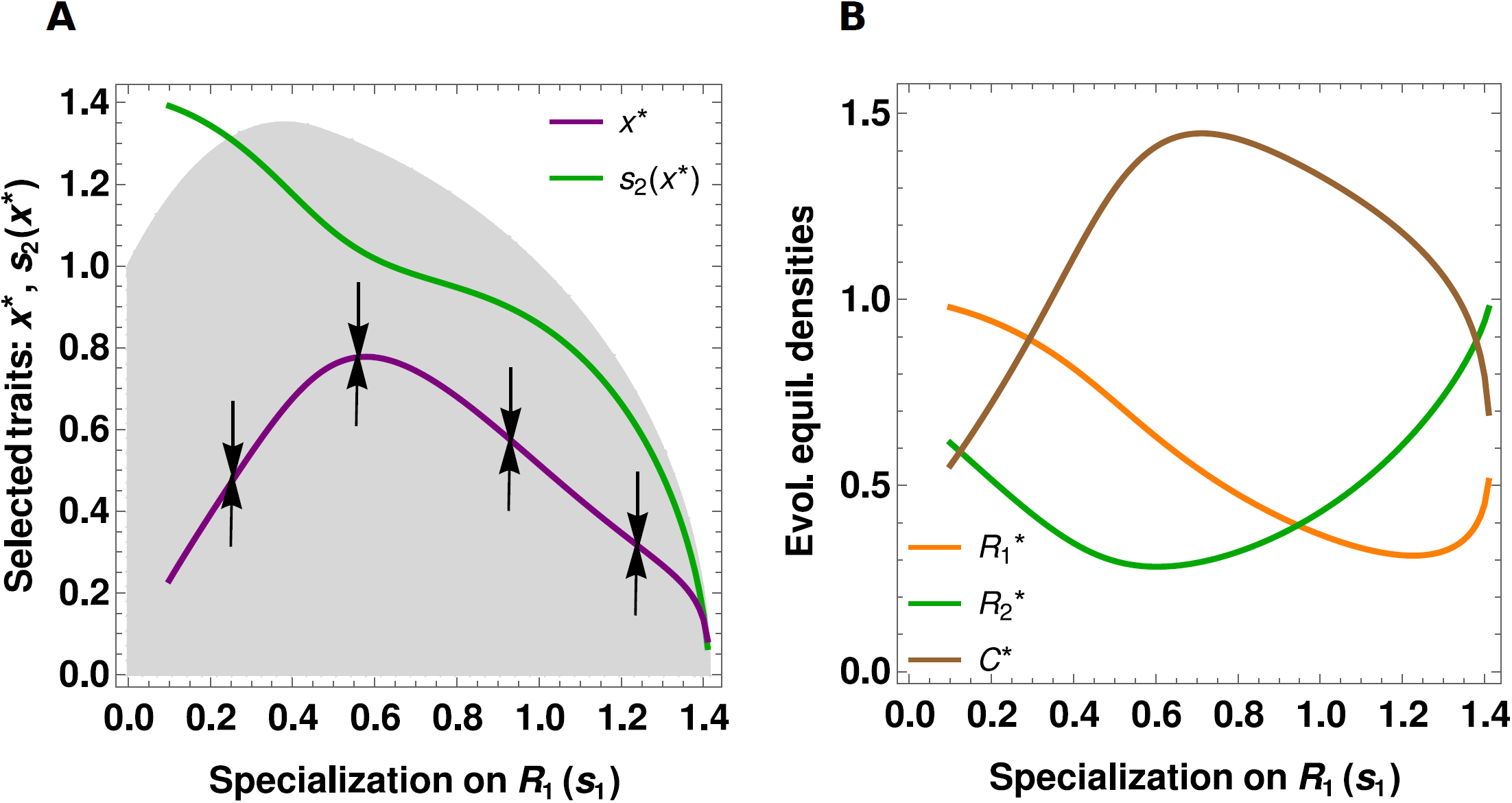
Effect of specialization on resource 1 on the evolutionary and ecological dynamics in the “opportunity cost” scenario. A) effect on the selected level of niche construction *x\** (in purple, the arrows indicate the direction of evolution) and subsequent consumption of resource 2, *s*_*2*_*(x*)* (in green). The area of stable ecological coexistence of the three species when varying agriculture and the specialization on *R*_*1*_ (i.e., in a *(s*_*1*_,*x)* plane) is represented in grey. B) effect on the species densities at the ecological evolutionary equilibrium (that is at the selected niche construction): the consumer is in brown, *R*_*1*_ is in orange, *R*_*2*_ is in green. *b*_*1*_ *= b*_*2*_ = 2, *g*_***1***_ *= g*_*2*_ *=* 2, *e*_*1*_ *= e*_*2*_ *=* 2, *L =* 2, *w =* 0.1, *m=*2, *k=*2, *u=*0.5

When specialization on the managed resource *s*_*1*_ is relatively low (left part of Fig 3 *A*), as predicted, increasing *s*_*1*_ leads to an increase in the selected agriculture intensity *x\**. The level of foraging on resource 2, *s*_*2*_*(x*)* consequently decreases due to trade-off constraints. However, when *s*_*1*_ is higher (right part of Fig 3 *A*), the pattern is reversed: increasing *s*_*1*_ selects for less agriculture. We can explain this by considering the evolutionary equilibrium densities (Fig 3 *B*): because an increase in *s*_*1*_ decreases the foraging on resource 2 (blue line, Fig 3 *A*), the alternative resource suffers less from the indirect effects. In the second part of the graph, it increases in density (green line). This increases its profitability which selects for less niche construction.

As in the “exploitation cost” scenario, the selected agriculture level (*x\**, purple line) falls in the coexistence area represented in gray for all specialization values. Evolution of agriculture therefore again favors the persistence of the system. This however depends on the trade-off shapes, and on the effect of niche construction on the helped resource, that mediates apparent competition and hence coexistence. We can obtain insight from how evolution could reduce coexistence from Fig 3 *B*: at low *s*_*1*_, increasing specialization on the helped resource decreases the alternative resource density at the evolutionary equilibrium. The evolution selects for more niche construction (advantaging its competitor) and higher consumer densities at the equilibrium, then the alternative resource suffers from a higher apparent competition. A even stronger pattern can lead to the extinction the alternative resource. When the helped resource obtains even higher benefits from niche construction (high conversion efficiency *w*), apparent competition is strong enough that for intermediate *R*_*1*_ specializations, *R*_*2*_ may become extinct (see Supplementary Figure S4). In this case, evolution kills the alternative resource but also destabilizes the community since it increases the agricultural trait to levels where the consumer and the helped resource are involved in a very strong positive feedback that produce an ever increasing dynamics.

#### Can evolution increase the diversity of the system?

We have seen that even though evolution mostly maintains the diversity in CSS scenarios (since the selected trait *x\** falls in the coexistence area) it can also disrupt it if apparent competition is too strong. We now investigate whether the diversity in the system can be increased, in particular through dimorphism in the consumer population.

In the “exploitation cost” scenario, because the non-invasibility and convergence criteria are equivalent, the two only types of evolutionary dynamics are evolutionary endpoints (CSS) that select for intermediate agriculture levels, or repellors leading to runaway evolution towards either no niche construction or full niche construction (potentially destabilizing for the community) or to intermediate levels of niche construction (see supplementary figure S1).

In the “opportunity cost”, all types of singular strategies can be potentially obtained, combining convergence and non-convergence, invasibility and non-invasibility. In particular, it is possible to obtain convergent points that are invasible: evolutionary branching points. In such conditions, the traits evolve towards the singularity value where disruptive selection is experienced. Dimorphism can then be maintained in the consumer population. We illustrate these dynamics in Figure 4. If the initial level of niche construction is high enough (that is, higher than the repellor value of the singularity, around 0.4 in the example), it evolves towards the singularity (around 1.2) where it experiences disruptive selection. We then observe the coexistence of two consumers in the population, a “farmer” strategy with a high value of niche construction investment and a “predator” strategy, not investing at all into niche construction. If initial niche construction investment is too low, it is then counter-selected and evolves towards zero (which is consistent with the positive nature of the feedbacks, we can expect threshold-dependent dynamics). This evolution leads to the coexistence of the two resources. The farmer strategy is a full specialist on the helped resource. The predator strategy consumes both preys, and is a relative generalist (here we have a higher specialization on the alternative resource, which is a property of our parameters: we fixed *s*_*1*_ to 0.5, so evolution towards zero niche construction automatically brings *s*_*2*_ to 1.5). The evolutionary branching point that we obtain here is not linked to the trade-off between specialization and niche construction shape (*k=*1) but to the niche “privatization” effect, that is the accelerating positive effect of agriculture intensity on the consumption efficiency on the resource (*u*=2). This is intuitive: providing diminishing returns to the farmer should favor generalism, while providing increasing returns favors specialization and more niche construction.

**Figure 4:**
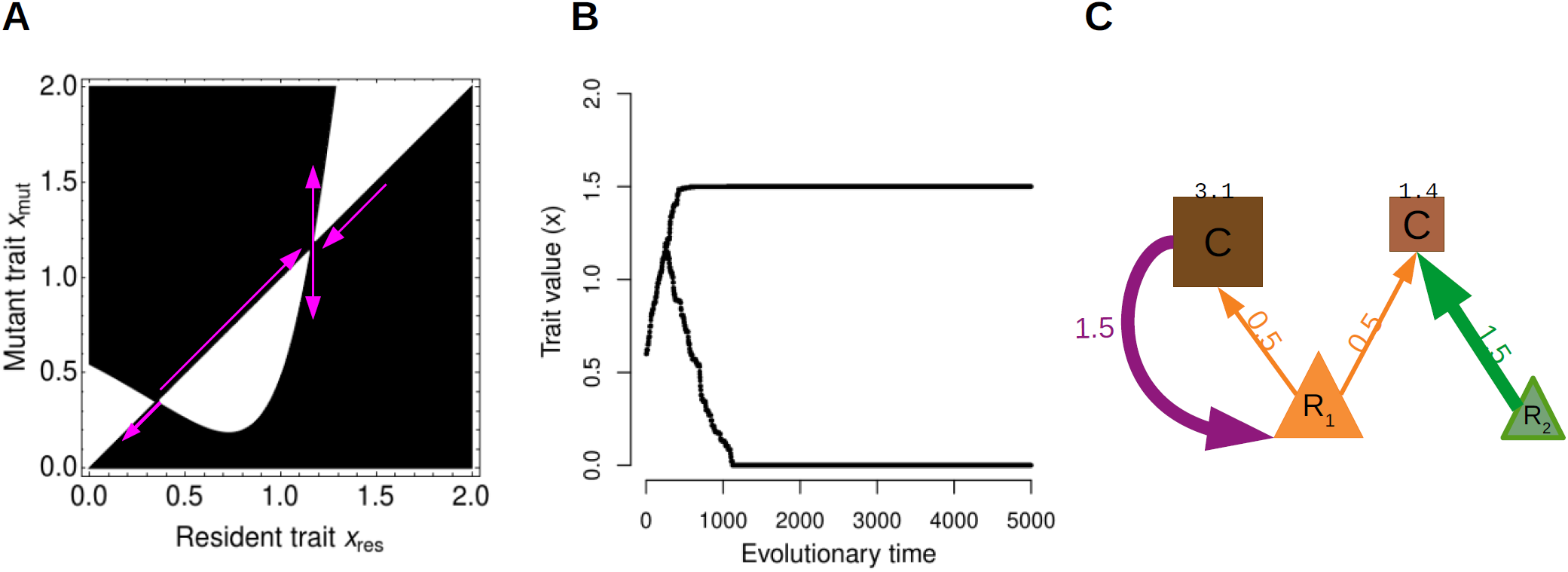
Evolutionary branching leading to dimorphism in agriculture investment, in the “opportunity cost” scenario. A: Pairwise Invasibility Plot, the sign of the fitness of a mutant of trait *x*_*mut*_ appearing in a population of trait *x*_*res*_ is shown (white for negative, black for positive). The pink arrows indicated the direction of evolution. B: Evolution trajectory of the investment into agriculture: the trait experiences disruptive selection at the convergent singularity leading to two coexisting phenotypes. C: final module with traits values (in color) and population densities (in black): the width of the arrows represents the trait value, not the net interaction (eg, the net per capita effect of *C* on *R*_*1*_ is *s*_*1*_ *-w x* = 0.35). Values are: *s*_*1*_*=0.5, k=1, u=2, w=0.1, b*_*1*_*=b*_*2*_*=3, g*_*1*_*=g*_*2*_*=2, e*_*1*_*=e*_*2*_*=2, m=2*.

### 3) Coevolution of the specialization and the agriculture investment traits

Until now we have fixed one trait and considered two trade-off scenarios, namely the opportunity cost and exploitation cost scenarios. We now assume that the three traits evolve simultaneously while still respecting the trade-off constraint, which means that we are looking at the coevolution of two traits, for instance *x* and *s*_*1*_, and with *s*_*2*_ being deduced from the two first and trade-off (eq (2)).

We investigate whether our prediction of a correlation between the profitability of the cultivated resource and the coevolution of specialization and agriculture on this resource. To do this, we vary the intrinsic growth rates of the two resources (*b*_*1*_ and *b*_*2*_) which can represent a proxy for their profitabilities, and explore how the values of the three traits at the coevolutionary equilibrium are impacted. We present these results in Figure 5. In Fig 5 *A*, *b*_*1*_ is varied and the values of the traits is plotted. We show that as the profitability of *R*_*1*_ increases, both *x\** and s_1_* increase, which means an increase in the consumer specialization on *R*_*1*_ and the investment in its cultivation. Inversely, when b_2_ is increased (Fig 5 *B*), the profitability of resource *R*_*2*_ increases, thus decreasing the evolution towards specialization and agriculture on *R*_*1*_. We then explore how the joint variation of the two profitabilities determine the ecological state at the coevolutionary equilibrium. We show how the selected agriculture investment (Fig 5 *C*) and relative preference on R_1_ (Fig 5 *D*) increase with increased relative profitability of *R*_*1*_ compared to *R*_*2*_. In Fig 5 *E*, we summarize the different possible states of the system with three areas. When *b*_*2*_ is high and *b*_*1*_ is low (area 1), the consumer is mostly specialized on the non cultivating resource at the equilibrium and investing less in constructing *R*_*1*._ When *b*_*1*_ is intermediate and *b*_*2*_ is high (area 2), the consumer is a generalist that cultivates *R*_*1*_ and consumes *R*_*1*_ and *R*_*2*_. When *b*_*1*_ is high, the consumer evolves to predominantly agricultural strategies with high investment into the cultivation and consumption of *R*_*1*_.

**Figure 5:**
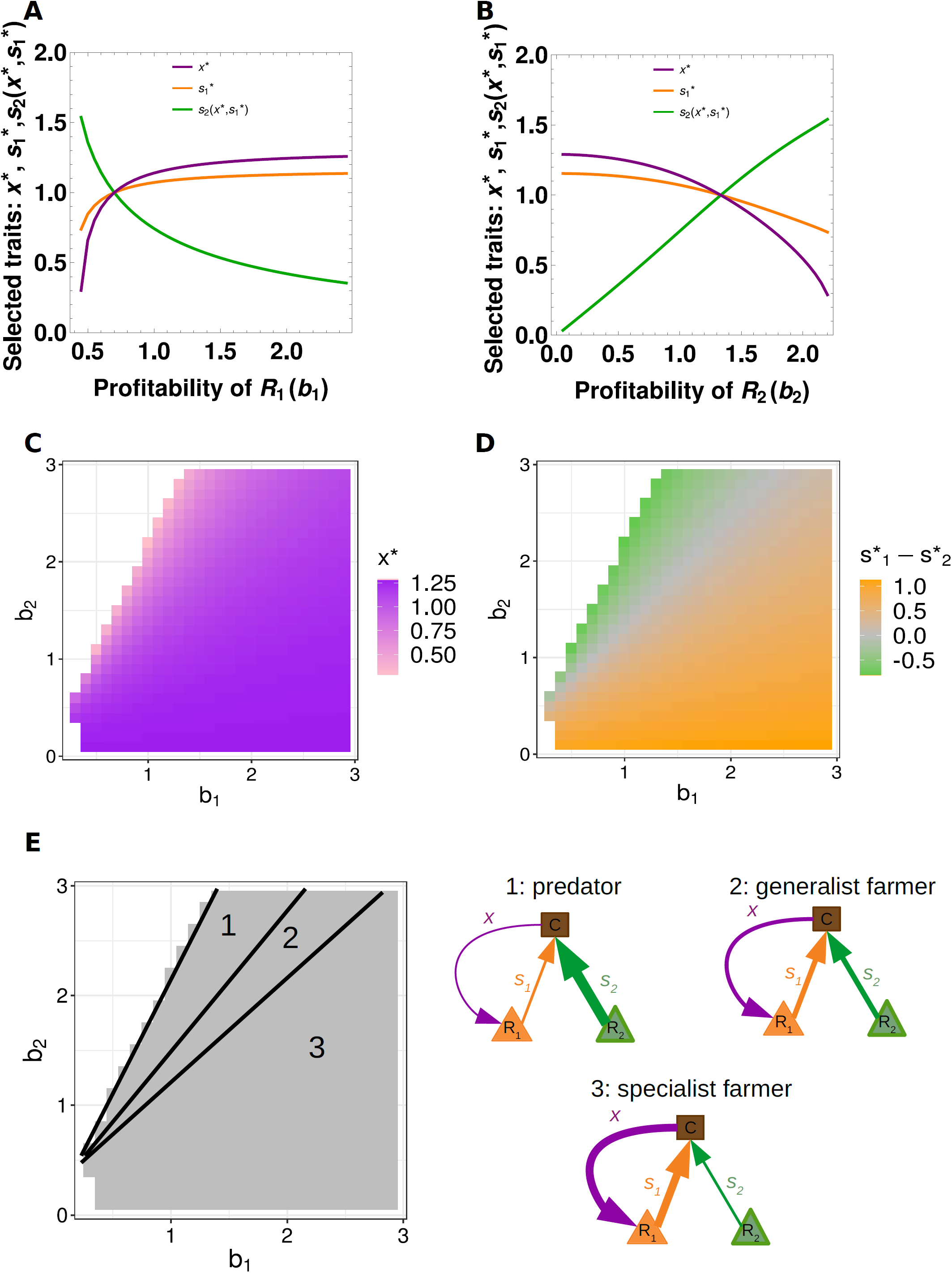
Coevolutionary dynamics. A) Correlation between the profitability of *R*_*1*_ and the values of the traits at the coevolutionary equilibrium (attractor, CSS): *x\** is shown in purple, *s*_*1*_*\** in orange, and *s*_*2*_*\** in green. B) Correlation between the profitability of *R*_*2*_ and the values of the traits at the coevolutionary equilibrium (attractor, CSS): *x\** is shown in purple, *s*_*1*_*\** in orange, and *s*_*2*_*\** in green. C) Values of the niche construction trait at the coevolutionary equilibrium in the two resources profitabilities plane. D) Relative resource preference at the coevolutionary equilibrium: the consumer is more specialized on *R*_*1*_ in the orange area and more specialized on *R*_*2*_ in the green area. E) Module state in the different areas defined by the profitabilities of the resources. When *b*_*2*_ is high and *b*_*1*_ is low (area 1), the consumer is mostly specialized on the non cultivating resource at the equilibrium and investing less in constructing *R*_*1*._ When *b*_*1*_ is intermediate and *b*_*2*_ is high (area 2), the consumer is a generalist that cultivates *R*_*1*_ and consumes *R*_*1*_ and *R*_*2*_. When *b*_*1*_ is high, the consumer evolves to predominantly agricultural strategies with high investment into the cultivation and consumption of *R*_*1*_. Values are: *g*_*1*_*=g*_*2*_*=*2, *e*_*1*_*=e*_*2*_*=*2, *L=*3, *w=*0.1, *k=*2, *u=*2, *m=*1. In panel A, *b*_*2*_=1. In panel B, *b*_*1*_ = 1.

## Discussion

In this work, we have studied the evolution of niche constructing agriculture in a consumer-resource perspective. Our initial prediction is that intuitively, cultivating a resource that the consumer is not consuming (so, a resource that is not profitable) is not adaptive and is counter-selected. We show a positive correlation between the evolution of agriculture and specialization on the cultivated resource, both in the two trade-off scenarios (although the cost of agriculture mitigates this result) and in the coevolution study.

In the “exploitation cost” scenario, we predicted that increasing the (fixed) specialization on the alternative resource should decrease the investment into agriculture. In the “opportunity cost” scenario, increasing the (fixed) specialization on the cultivated resource should increase the investment into agriculture. These predictions are partly confirmed by our results. Indeed, increasing the profitability of the alternative resource (via increased specialization) decreases the investment for agriculture in the “exploitation cost” scenario. Conversely, increasing the specialization on the cultivated resource increases the selected level of agriculture in the “opportunity cost” scenario, although a further increase mitigate the profitability of the cultivated resource through indirect effects, leading to a decrease in the selected level of agriculture.

The joint evolution of specialization and agriculture is analyzed in the coevolution scenarios, where an increase in the profitability of the cultivated resource leads to an increase in the level of investment into both specialization of this resource and cultivation of the resource, via the emergence of an eco-evolutionary feedback between these two traits. This gives important insights on the transitions from foraging-only to agriculture-only strategies, that may be observed in the model, as well as on transitions from foraging-only to facultative farmers (or facultative foragers).

This question that we here investigated here from a niche construction/agriculture perspective is well suited to understand the evolution of mutualism or symbiosis. For instance, some ants perform facultative agriculture (on aphids or fungi) and usually remain generalist, while other taxa are obligate farmers of fungi (Chapela et al. 1994; Stadler and Dixon 2005) or aphids (Ivens et al. 2016) in which cases many specialized traits are observed. We here show that such dynamics may emerge from foraging theory constraints (that may be linked to physiology, but also to resource densities, dynamics). In our model, we do not investigate whether the net interaction is mutualistic for the helped resource, because this has been proven difficult to measure experimentally (context-dependency of interactions (Chamberlain et al. 2014), multiple components of fitness) but we implement both a positive and a negative aspect of the interaction. For instance, in the ant-aphid interaction, it is unclear whether this can be qualified as mutualism or exploitation (Offenberg 2001; Stadler and Dixon 2005; Billick et al. 2007).

One of the two questions that stimulated our study was to improve our understanding of the conditions under which agriculture may evolve. The other one, is the effect of this evolution on the ecological dynamics and in particular the coexistence of cultivated and non-cultivated resources. This aspect is here mediated by the indirect interactions occurring in the system, particularly apparent competition. Apparent competition has been experimentally described in agricultural systems where a predator has a positive effect of one of its preys, notably in aphid-tending ant increasing the predation on other arthropods (Warrington and Whittaker 1985; Wimp and Whitham 2001). In our model, we note that evolution does not optimize densities: this result that could seem counter-intuitive is rather well known when considering adaptive dynamics studies (Metz et al. 2008; Boudsocq et al. 2011). Second, we show that evolution of agriculture may favor coexistence in the system. In the “exploitation cost” scenario, without evolution, we expect that increasing specialization on the alternative resource should reduce coexistence, as it increases predation pressure on the resource which does not benefit from niche construction. When we consider evolution though, the decreased selected level of niche construction compensates for this effect and rescues the alternative resource, thus favoring coexistence. We can observe similar patterns in the “opportunity cost” scenario. Adaptive foraging is also known to improve coexistence of resources by mitigating apparent competition in purely trophic systems (Abrams 2010). We note however that this evolutionary facilitation of coexistence is dependent on costs and benefits. Especially, in cases where the efficiency of niche construction is very high, even if the evolution mitigates the stronger apparent competition that the alternative resource suffers from, it might not be enough to rescue it. Such effects may be linked to natural observations. In devil’s gardens, ants kill competitors of their host plant species thereby actively limiting coexistence at a local scale (Frederickson et al. 2005). In our model, runaway evolution toward high niche construction may yield large positive feedbacks, potentially destabilizing the system and leading to the exponential growth of the consumer and helped resource. Such infinite increases could be avoided if we considered diminishing returns of niche construction, for instance, if niche construction is limited by other parameters. This addition to the model has been shown to stabilize the ecological dynamics within this module (Picot et al. 2019).

Evolution of agriculture in our model assumed two main hypotheses. First, that agriculture occurs at a cost, here constraining the foraging of resources (either the cultivated one or the alternative one) ; second, that agriculture provides a direct benefit to the organism that performs it. This direct benefit can also be interpreted as a trade-off between the consumption efficiency of resource 1 and its non-cultivation, and allows the evolution of agriculture because it limits the invasion of cheaters that would not pay the cost of agriculture investment. Such a scenario has been described in a model of niche construction in which pleiotropy allows the maintenance of niche construction (Chisholm et al. 2018). If there is a positive correlation between the investment and the returns, costly niche construction may evolve. If the direct benefits of niche construction are accelerating with increasing niche construction intensity, evolution can lead to dimorphism in the consumer population. We then observe the coalition of a highly specialized farmer, that invests a lot in niche construction and only preys on the helped resource, and of a generalist predator strategy that performs no or very low agriculture. This latter could be considered a cheater in the evolution of cooperation terminology: it consumes the resource that is maintained by niche construction, without paying the cost of agriculture (but see Ghoul et al. (2013). Because of the direct benefit that the farmer obtains from the exploitation of the helped resource (“privatization” or “monopoly” (Krakauer et al. 2009; Chisholm et al. 2018; Scheiner et al. 2021), the invasion of the cheater is avoided and the two phenotypes coexist.

Increasing the number of consumers also stabilizes the system: similar to exploitative competition where two consumers may coexist when consuming two shared resources (according to the *R*\* rule, (Tilman 1980)), in apparent competition, resources are more likely to coexist if they are consumed by *n* consumers because it balances apparent competition (similarly to the *P\** rule, (Holt et al. 1994)). If the consumer population is considered as a whole comprised of two subpopulations, we can then interpret this coalition as division of labor, or functional specialization, between farmers and foragers. This type of division of labour has been investigated in theoretical works which stress the importance of considering high benefits to specialization or accelerating returns (Rueffler et al. 2012; Cooper and West 2018). This is consistent with our model, and recalls the importance of trade-off geometries in evolutionary dynamics of specialization (de Mazancourt and Dieckmann 2004; Egas et al. 2004; Ravigné et al. 2009; Kisdi 2014). From an evolution of specialization perspective, the coalition between niche-constructors and non-constructors is also a coalition between a specialist and a generalist. The question of the coexistence of specialists and generalists have often been frequently investigated from a purely trophic context (Egas et al. 2004; Abrams 2006; Rueffler et al. 2006). Our model provides various dynamics that emerge from simple assumptions, and these results can be understood both in the evolution of cooperation and evolution of specialization framework. The coevolution analysis allows to describe situations where if a potentially cultivable resource is profitable enough, it might be advantageous to switch from a generalist forager to a specialized farmer, which can explain evolutionary transitions from foraging-only to settled agriculture behaviours.

