## Supplementary Information for "Implications of (co)evolution of agriculture and resource foraging for the maintenance of species diversity and community structure"

### 1) Mathematical analysis: details

#### “Exploitation cost” scenario: convergence and invasibility

The convergence and invasibility properties are expressed as functions of second derivatives of fitness. The singularity is non-invasible (evolutionary stable) if:

$$\left.\frac{\partial^2 W(x_{mut}, x_{res})}{\partial x_{mut}^2}\right|_{x_{mut} \to x_{res} \to x^*} < 0 \Leftrightarrow \quad (1)$$
$$e_1''(x^*) s_1(x^*) + 2 e_1'(x^*) s_1'(x^*) + e_1(x^*) s_1''(x^*) < 0$$

The term $2 e_1'(x^*) s_1'(x^*)$ is negative. Hence, the non-invasibility condition is dependent on trade-off concavity between *x* et $s_1$ ($s_1''(x^*_t)$ ), as well as on the benefit from niche construction ($e_1''(x^*)$ ). The singularity is convergent if:

$$\left.\frac{\partial^2 W(x_{mut}, x_{res})}{\partial x_{mut}^2}\right|_{x_{mut} \to x_{res} \to x^*} + \left.\frac{\partial^2 W(x_{mut}, x_{res})}{\partial x_{mut} \partial x_{res}}\right|_{x_{mut} \to x_{res} \to x^*} < 0 \quad (2)$$

Given that $\frac{\partial^2 W(x_{mut}, x_{res})}{\partial x_{mut} \partial x_{res}} = \left(e_1'(x_{mut}) s_1(x_{mut}) + e_1(x_{mut}) s_1'(x_{mut})\right) \frac{\partial R_1^*(x_{res})}{\partial x_{res}}$ , we have

$$\left.\frac{\partial^2 W(x_{mut}, x_{res})}{\partial x_{mut} \partial x_{res}}\right|_{x_{mut} \to x_{res} \to x^*} = \left(e_1'(x^*) s_1(x^*) + e_1(x^*) s_1'(x^*)\right) \left.\frac{\partial R_1^*(x_{res})}{\partial x_{res}}\right|_{x_{mut} \to x_{res} \to x^*} = 0 \quad (3)$$

Hence we conclude that the singularity is convergent if and only if it is non-invasible.

### “Opportunity cost” scenario: convergence and invasibility

The strategy is non-invasible if:

$$e_1''(x^*)s_1 R_1^*(x^*)+e_2 s_2''(x^*)R_2^*(x^*)<0 \tag{4}$$

And it is convergent if:

$$\begin{aligned}&e_1''(x^*)s_1 R_1^*(x^*)+e_2 s_2''(x^*)R_2^*(x^*)\\&+e_1'(x^*)s_1 \left.\frac{dR_1^*(x_{res})}{dx_{res}}\right|_{x_{res}\to x^*}+e_2 s_2'(x^*)\left.\frac{dR_2^*(x_{res})}{dx_{res}}\right|_{x_{res}\to x^*}<0\end{aligned} \tag{5}$$

With the trade-off functions that we considered, the fitness gradient at the singularity becomes:

$$G(x^*,x^*)=\left.\frac{\partial W(x_{mut},x_{res})}{\partial x_{mut}}\right|_{x_{mut}\to x_{res}\to x^*}=e_1 u x^{*u-1}s_1 R_1^*(x^*)-e_2 x^{*\frac{1}{k}-1}(L-x^{*k}-s_1^k)^{\frac{1}{k}-1}R_2^*(x^*)$$

We note that in this scenario, the resource densities at the ecological equilibrium can have an effect on the dynamics. In the linear case, the singular strategy cannot be studied (degenerate case with the nulling of the invasibility and convergence properties).

The non-invasibility condition becomes:

$$e_1(u-1)u x^{u-2}s_1 R_1^*(x^*)+e_2(1-k)(L-s_1^k)x^{k-2}(L-x^{*k}-s_1^k)^{\frac{1}{k}-2}R_2^*(x^*)<0$$

We get a sufficient condition for non-invasibility: $u < 1$ and $k > 1$. A sufficient condition for the invasibility is that if $u>1$ and $k<1$.

### 2) Supplementary Figures

#### “Exploitation cost” scenario

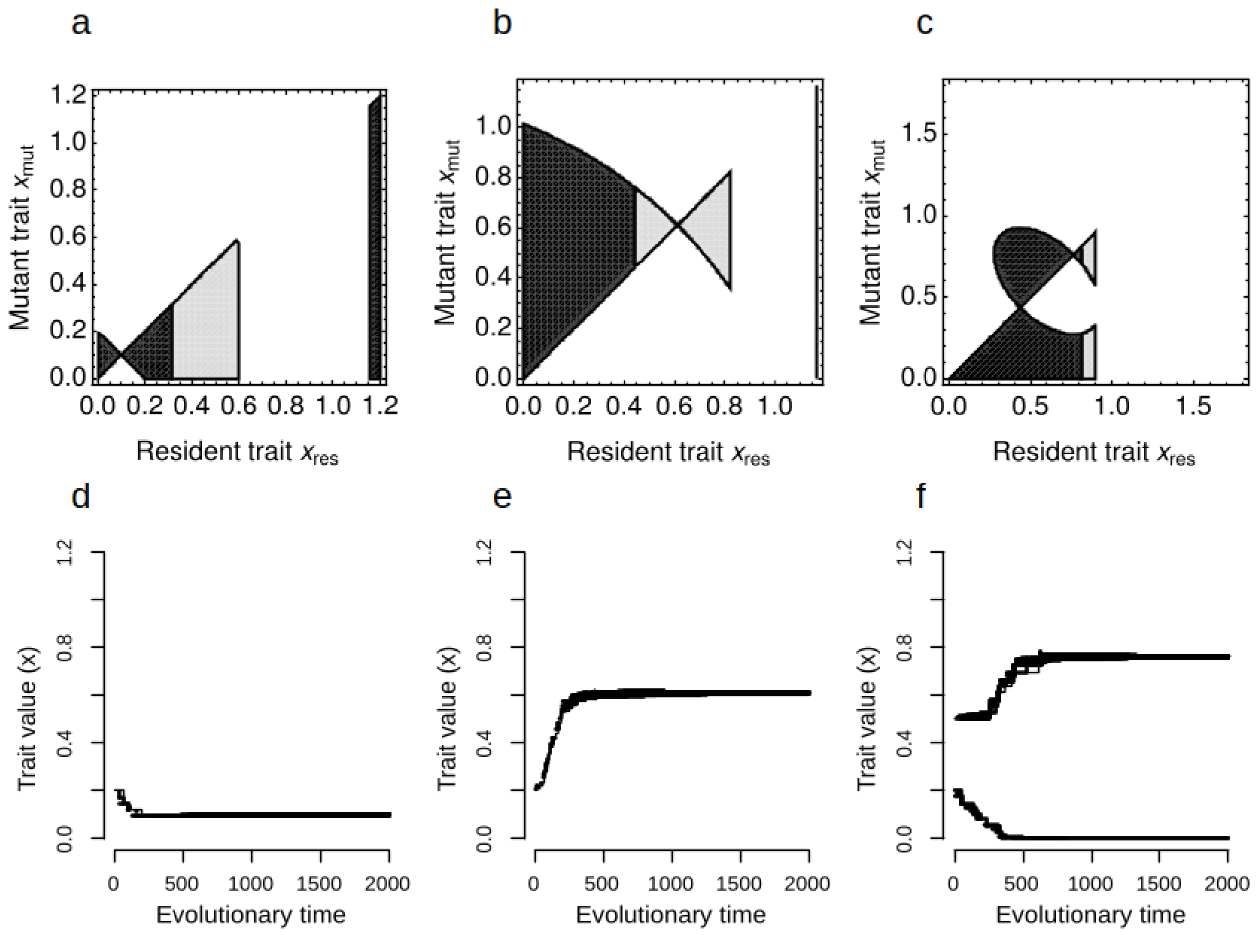


*Figure S1: Pairwise Invasibility Plot and corresponding evolutionary trajectories in the “exploitation cost” scenario. For panels a,b,c: the sign of the fitness of a mutant of trait $x_{mut}$ appearing in a population of trait $x_{res}$ is shown (white for negative, black and grey for positive). In the black area, the three species coexist stably, while in the gray area only the constructor and $R_1$ coexist. a: k=2, u=1, $s_2$=0.8 (CSS, evolutionary attractor) illustrated in d. b: k=1, u=1, $s_2$=0.8 (CSS), illustrated in e. c: k=1, u=2, $s_2$=0.2 (repellor and CSS, evolutionary bistability, illustrated in f). Other parameter values are $b_1 = b_2 = 2$, $g_1 = g_2 = 0.8$, $e_1 = e_2 = 1$, L =2, w = 1, m = 0.8.*

On panel 3a,b we obtain a CSS: the evolution leads the trait to an intermediate level of niche construction. On panel 3c, there is a repellor and a CSS: that is starting from less than 0.5 (the position of the repellor), evolution leads to zero niche construction, while starting from above 0.5, the evolution leads to the CSS that is an intermediate value of niche construction (0.8). The evolution brings the trait to a coexistent zone on panel 3b,c while it reduces diversity by killing resource 2 on panel a.

We present the effect of specialization on the selected level of agriculture when varying the weight of each trait in the trade-off function:

$$\gamma x^k + \beta s_1^k + \alpha s_2^k = L \quad (6)$$

This leads to altered distribution of the cost of increased specialization on $R_2$ between the selected agriculture level and specialization on $R_1$ (figure S2).

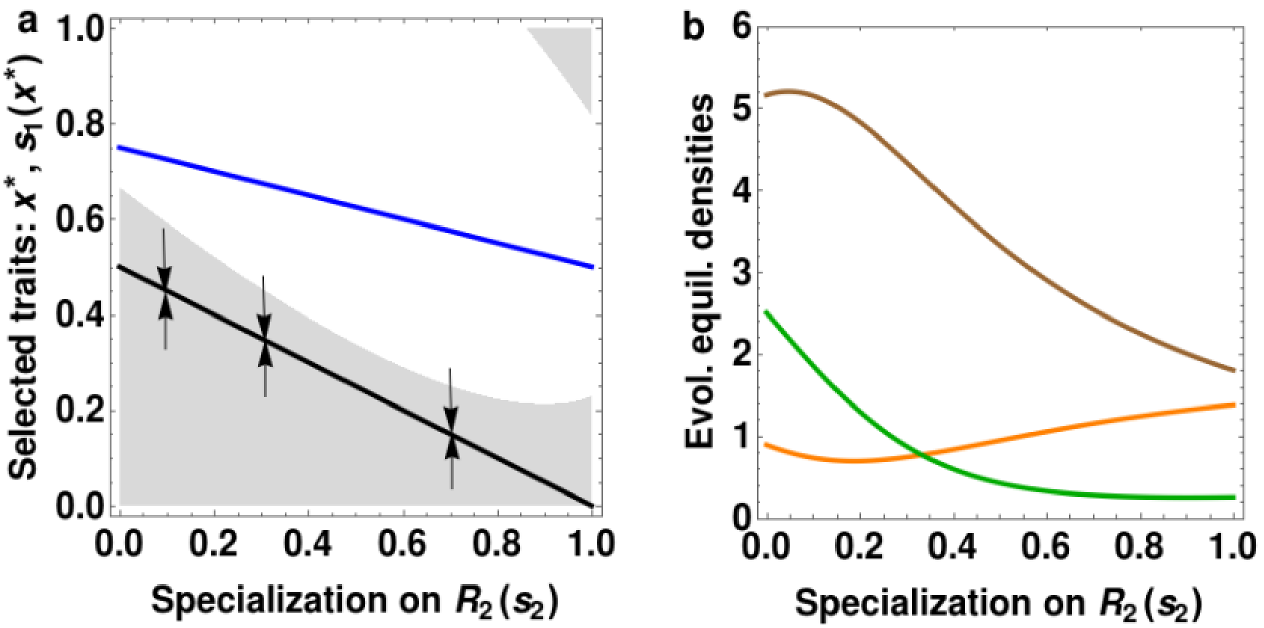


*Figure S2: Effect of specialization on resource 2 on the evolutionary and ecological dynamics in the "exploitation cost" scenario. a) effect on the selected level of niche construction x* (in black) and subsequent consumption of resource 2, $s_1(x^*)$ (in blue). The area of stable ecological coexistence of the three species when varying agriculture and the specialization on $R_2$ (ie, in a ($s_2$,x) plane) is represented in grey. b) effect on the species densities at the ecological evolutionary*

*equilibrium (that is at the selected niche construction): the consumer is in brown, $R_1$ is in orange, $R_2$ is in green. $b_1 = b_2 = 2$, $g_1 = g_2 = 0.8$, $e_1 = e_2 = 1$, $L = 2$, $w = 1$, $m = 0.8$, $k = 1$, $u = 1$, $\alpha = 1$, $\gamma = 1$, $\beta = 2$*

When the strategy is a repellor, we obtain a positive correlation between its value and the specialization on $R_2$ (figure S3): this leads to a negative correlation in terms of outcome of evolution, because the higher the specialization on $R_2$, the higher the repellor singular strategy, and the lower the range of initial values that lead to a high niche construction.

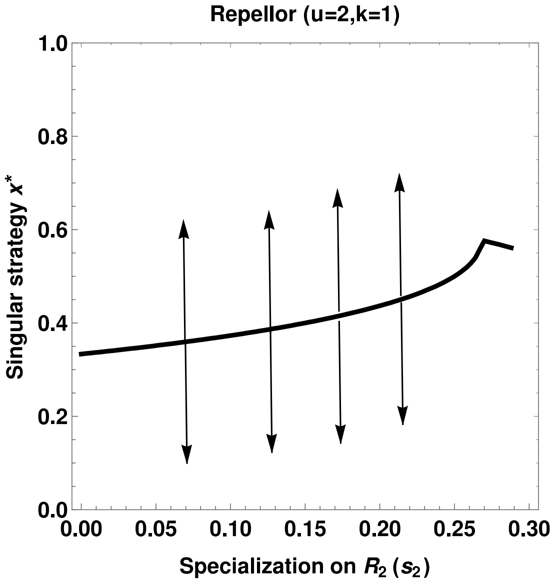


*Figure S3: The effect of specialization on resource 2 on the evolutionary dynamics. We plot the value of the repellor singular strategy (unstable evolutionary equilibrium) against the value of $s_2$. $b_1 = b_2 = 2$, $g_1 = g_2 = 0.8$, $e_1 = e_2 = = 1$, $L = 2$, $w = 1$, $m = 0.8$, $k = 1$, $u = 2$*

### “Opportunity cost” scenario

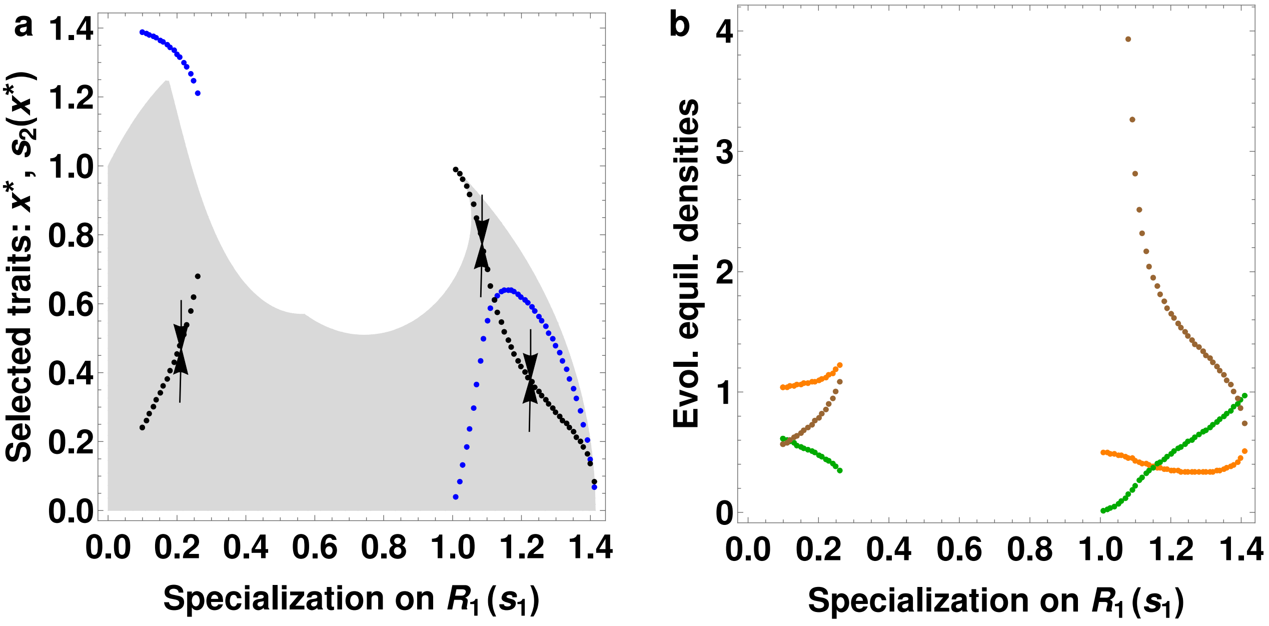


*Figure S4: Effect of specialization on resource 1 on the evolutionary and ecological dynamics in the “opportunity cost” scenario. a) effect on the selected level of niche construction x* (in black) and subsequent consumption of resource 2, $s_2(x^*)$ (in blue). The area of stable ecological coexistence of the three species when varying agriculture and the specialization on $R_1$ (ie, in a $(s_1,x)$ plane) is represented in grey. b) effect on the species densities at the ecological evolutionary equilibrium (that is at the selected niche construction): the consumer is in brown, $R_1$ is in orange, $R_2$ is in green. $b_1 = b_2 = 2$, $g_1 = g_2 = 2$, $e_1 = e_2 = 2$,, $L = 2$, $w = 1$, $m=2$, $k=2$, $u=0.5$*
